# PPAD: A deep learning architecture to predict progression of Alzheimer’s disease

**DOI:** 10.1101/2023.01.28.526045

**Authors:** Mohammad Al Olaimat, Jared Martinez, Fahad Saeed, Serdar Bozdag, Alzheimer's Disease Neuroimaging Initiative

## Abstract

Alzheimer’s disease (AD) is a neurodegenerative disease that affects millions of people worldwide. Mild cognitive impairment (MCI) is an intermediary stage between cognitively normal (CN) state and AD. Not all people who have MCI convert to AD. The diagnosis of AD is made after significant symptoms of dementia such as short-term memory loss are already present. Since AD is currently an irreversible disease, diagnosis at the onset of disease brings a huge burden on patients, their caregivers, and the healthcare sector. Thus, there is a crucial need to develop methods for the early prediction AD for patients who have MCI. Recurrent Neural Networks (RNN) have been successfully used to handle Electronic Health Records (EHR) for predicting conversion from MCI to AD. However, RNN ignores irregular time intervals between successive events which occurs common in EHR data. In this study, we propose two deep learning architectures based on RNN, namely Predicting Progression of Alzheimer’s Disease (PPAD) and PPAD-Autoencoder (PPAD-AE). PPAD and PPAD-AE are designed for early predicting conversion from MCI to AD at the next visit and multiple visits ahead for patients, respectively. To minimize the effect of the irregular time intervals between visits, we propose using age in each visit as an indicator of time change between successive visits. Our experimental results conducted on Alzheimer’s Disease Neuroimaging Initiative (ADNI) and National Alzheimer’s Coordinating Center (NACC) datasets showed that our proposed models outperformed all baseline models for most prediction scenarios in terms of F2 and sensitivity. We also observed that the age feature was one of top features and was able to address irregular time interval problem.

## 1 Introduction

Alzheimer’s Disease (AD) is an irreversible neurodegenerative disease that leads to problems in cognitive functioning (e.g., memory loss and impaired reasoning) and behavioral changes (e.g., aggression, wandering, and anxiety). Unusual accumulation of amyloid plaques and neurofibrillary tangles in the brain are considered as the main causes of AD (Lee et al., 2019). According to the World Health Organization (WHO), there are about 40 million AD cases worldwide. In the United States, there are about 6 million AD cases, and this number is expected to reach 14 million by 2050 (Venugopalan et al., 2021; Alzheimer’s Association, 2015).

Mild cognitive impairment (MCI) is an intermediary stage between cognitively normal (CN) state and AD. MCI is determined through an impairment of memory on standard tests with the absence of significant impairment in daily living activities and dementia (Winblad et al., 2004). Using a standardized test, impairment on cognitive is defined as performance below 1.5 standard deviations (SD) of the age-, sex- and education-adjusted normative mean; according to the test, MCI can be classified based on the severity into early and late MCI. Early MCI (EMCI) refers to a case when the performance is between 1.0 SD and 1.5 SD below the normative mean on the test, whereas late MCI (LMCI) refers to a case when the performance is 1.5 SD below the normative mean on the test (Jessen et al., 2014; Aisen et al., 2010). About 15% of MCI patients convert to AD per year while 80% of MCI patients convert to AD within about six years (Tábuas et al., 2016). MCI cases who progress to AD eventually are called MCI-converter and MCI cases who stay as MCI or revert to CN are called MCI-non-converter.

The diagnosis of AD can be made after significant symptoms of dementia such as short-term memory loss are already present. The diagnosis after the onset of the disease creates emotional burden for patients and their family members and economic burden to the healthcare. The estimated healthcare cost for AD was over $300 billion in 2020 (Wong et al., 2020). As a result, developing a robust method that can early predict conversion from MCI to AD is crucial for patients to have better treatments, interventions to delay or prevent AD progression.

Electronic Health Record (EHR) is a sequential data represented as temporal sequences of clinical features and has been used to train machine learning models to classify and cluster patient records for improving clinical decision making. Traditional machine learning methods, however, aggregate clinical features; thus ignores temporal relations between these sequences. Recurrent Neural Network (RNN) is a deep learning model used to process sequential data and maintains temporal relations between sequences (Goodfellow et al., 2016). Long Short-Term Memory (LSTM) and Gated Recurrent Unit (GRU) are RNN variants that have the capability to handle long-term dependencies, which is considered the drawback of vanilla RNN. The main difference between the GRU and LSTM is their complexity (i.e., number of learnable parameters). The GRU is a less complex architecture than LSTM, thus are more preferable especially when training data is not abundant (Greff et al., 2016).

To identify biomarkers of conversion from MCI to AD, various machine learning methods have been utilized. In (Zhang et al., 2012), support vector machine (SVM) was applied with a multi-task learning to identify AD biomarkers and predict the 2-years conversion from MCI to AD using baseline measurements from Magnetic Resonance Imaging (MRI) and Cerebrospinal fluid (CSF). The proposed model achieved 73.9% accuracy, 68.6% sensitivity, and 73.6% specificity. In (Cheng et al., 2015), a domain transfer learning model was proposed for using not only MCI samples but also AD and CN as auxiliary samples to identify biomarkers that can be used for a classification task to distinguish between MCI-converter and MCI-non-converter samples. The proposed model achieved 79.4% accuracy, 84.5% sensitivity, and 72.7% specificity.

Integration of multi-modality data has been performed to improve the performance of predicting conversion from MCI to AD by extracting AD biomarkers from each modality. A graph-based semi-supervised learning method that integrates brain image data from MRI and Positron Emission Tomography (PET) was proposed to distinguish between EMCI from CN cases (Kim et al., 2013). The proposed method achieved 68.5% accuracy, 53.4% sensitivity, and 77% specificity. In (Lee et al., 2019), a multi-modal GRU model was trained using longitudinal cognitive performance and CSF biomarkers data, and cross-sectional neuroimaging and demographic data to predict MCI to AD conversion. The results showed that the proposed model achieved 81% accuracy and an area under the receiver operation characteristics curve (AUC) of 86%. In (Venugopalan et al., 2021), an integrative classification method was proposed to classify patients into AD, MCI, and CN. The model was trained on clinical and genetic features extracted using stacked denoising auto-encoders and brain image features extracted using Convolutional Neural Network (CNN). The results have shown that integrating multi-modality data outperforms single modality models.

In (Nguyen et al., 2020), an RNN model was applied to the longitudinal cognitive performance, MRI, and CFS data af 1677 samples in Alzheimer’s Disease Neuroimaging Initiative (ADNI) database to predict the diagnosis of patients in the future up to six years and achieved better performance than baseline models. In (Li et al., 2019), a deep learning model based on an LSTM autoencoder was proposed to extract the hidden temporal pattern in longitudinal data for five cognitive measures for one-year follow-up. The new extracted features were combined with baseline hippocampal measures extracted from MRI scans to train and evaluate a prognostic model using Cox regression to predict AD progression for MCI individuals.

In analyzing longitudinal biomedical data, irregular time intervals between clinical visits poses a technical challenge. Deep learning methods that can handle sequential data (e.g., RNN) assume that equal intervals between inputs in the sequence. To address this issue, Time-Aware LSTM (T-LSTM) was proposed to modify the memory state the current cell state based on the time gap between the current and previous cell states (Baytas et al., 2017). The results on progression of Parkinson’s disease data showed improved performance than baseline methods. In another study, T-LSTM was evaluated on synthetic and real data for Chronic Kidney Disease (Luong et al., 2018). The results showed that T-LSTM autoencoder can be used to deal with sequential data to generate the latent space of the longitudinal profile, but the latent space of the longitudinal of real data was not able to subtype Chronic Kidney Disease.

Studies have shown an association between AD and several genes such as CTNNA3, GAB2, PVRL2, and TOMM40. Among these, epsilon4 allele of apolipoprotein E gene (APOEɛ4) is the most important genetic risk factor for AD (Chalmers et al., 2003). In addition, demographics such as age, gender, alcohol consumption, smoking, depression, head injury, education, race, ethnicity, and nutrition have also been reported as nongenetic risk factors (Ikeda et al., 2010; Hall et al., 1998). As a result, these genetic and nongenetic risk factors can play role if they are utilized by a predictive model for AD progression.

In general, irregular number of visits for patients, irregular intervals between visits, and missing values are common drawbacks of EHR. Since datasets from AD cohorts suffer from the same drawbacks, there is a crucial need to develop methods for early predicting conversion from MCI to AD while addressing such data irregularities. Most of existing methods do not consider the irregular time intervals between consecutive visits and give equal weight to sensitivity and specific of the model. However, increasing sensitivity (i.e., correctly predicting individuals who would convert to AD) is more important. Moreover, several existing studies do not integrate longitudinal data with cross-sectional demographics data such as gender, race, ethnicity, patients’ education, and APOEɛ4. Most tools are also not made publicly availability, which limits their application to new datasets.

In this study, we propose two open-source deep learning models, PPAD and PAD-AE, for early predicting conversion from MCI to AD at the next visit and multiple visits ahead for patients, respectively. PPAD and PPAD-AE integrate longitudinal features with cross-sectional demographic data. To minimize the effect of the irregular time intervals between visits, we propose using age in each visit as an indicator of time change between consecutive visits. In addition, we utilized a customized loss function to give more weight on predicting conversion to AD cases, thereby increasing the model’s sensitivity. To show robustness of our proposed models, we used two evaluation setups, by which (i) ADNI dataset was used to train and test proposed models; (ii) ADNI dataset was used to train proposed models and the National Alzheimer’s Coordinating Center (NACC) dataset was used to test proposed models. Our experimental results showed that our proposed models outperformed all baseline models for most of the prediction scenarios in terms of F2 and sensitivity. We also demonstrated that using age feature improved the model performance by helping address the irregular time interval between consecutive visits. We made PPAD and PPAD-AE publicly available at https://github.com/bozdaglab/PPAD under Creative Commons Attribution Non-Commercial 4.0 International Public License.

## 2 Materials and Methods

### 2.1 Datasets

In this study, longitudinal and cross-sectional data from two datasets were used. The main dataset was ADNI database (adni.loni.usc.edu). The ADNI was launched in 2003 as a public-private partnership. The primary goal of ADNI has been to test whether serial magnetic resonance imaging (MRI), positron emission tomography (PET), other biological markers, and clinical and neuropsychological assessment can be combined to measure the progression of mild MCI and early AD. Since it has been launched, the public-private cooperation has contributed to significant achievements in AD research by sharing data to researchers from all around the world (Jack et al., 2010; Jagust et al., 2010; Saykin et al., 2010; Trojanowski et al., 2010; Weiner et al., 2010; Risacher et al., 2010; Weiner et al., 2013).

The second dataset was NACC (Besser et al., 2018), which is a large, centralized resource for AD research. It contains data from multiple study sites across the United States, including demographic, cognitive, genetic, and MRI data. The database is designed to facilitate research on the causes, diagnosis, and treatment of AD.

In this study, we have used two different experiment setups to train and evaluate our proposed models. In the first setup, ADNI dataset was split into training and test datasets to train and test the proposed models. In the second setup, the whole ADNI dataset was used to train the proposed models while NACC dataset was used as an external dataset to test the proposed models.

#### 2.1.1 ADNI dataset for training and testing the proposed models

To obtain longitudinal and cross-sectional data from all ADNI studies (ADNI1, ADNI2, and ADNI-GO), ADNImerge R package was used (https://adni.bitbucket.io/) (McCombe et al., 2020; Jiang et al., 2020). The raw data consisted of 15,087 records from 2288 unique patients. Each record represents a patient visit that consists of feature values from four data modalities namely cognitive performance measurement, MRI, CSF, and demographic, and the diagnosis label (i.e., CN, MCI or AD). The dataset had several irregularities: the patients had varying number of visits ranging from 1 to 21; the time between consecutive visits for a patient varied from 3 months to 60 months; and several visits had missing feature values. To address these issues, we preprocessed the dataset by performing following steps: First, irrelevant features such as name of ADNI project and duplicated columns (i.e., the feature that had the same value of another feature with a different name) were removed. In total, 55 such features were discarded. Second, all visits with missing values rate >40% were discarded. In total, 4979 visits were discarded. Third, 128 patients who do not have APOEɛ4 status were removed resulted in discarding 154 visits. Fourth, all visits with CN diagnosis were discarded since our goal was to analyze progression from MCI to AD. We also discarded patients who had only a single visit as it provided no disease progression information. In total, 4195 visits were discarded. Finally, we discarded 32 features that had missing values in >60% of the records.

For the remaining features that had missing values, K Nearest Neighbor (KNN) algorithm was employed to impute the missing values. Each missing feature was imputed using the average of values from nearest neighbors that had a value for that feature. When finding the nearest neighbors, only the records with the same diagnosis (i.e., MCI or AD) were considered. The Euclidean distance metric was used and the number of neighbors, *k*, was set to five. After imputation, the final dataset had 20 longitudinal and five demographics features for 1169 patients and 5759 visits. To train the proposed models, the dataset was randomly split in a stratified manner into 70% train and 30% test data. After splitting, each feature was normalized using min-max normalization. For generalization, the whole procedure was repeated for three random splits.

#### 2.1.2 ADNI dataset for training and NACC dataset for testing the proposed models

In this setup, we performed data harmonization to collect the overlapping features from ADNI and NACC datasets. To do so, first, we obtained three different data modalities from NACC dataset namely cognitive, genetic, and MRI data. Demographic information including gender, race, ethnicity, and education were in the cognitive data. We obtained APOEɛ4 status from genetic data for all the patients in the cognitive data. We could not integrate MRI data features with cognitive data due to the low number of overlapping patients, mismatching number and date of visits for overlapping patients, and high rate of missing values (> 90%) of overlapping features between ADNI and NACC datasets. At the end, we harmonized nine features between ADNI and NACC datasets (Supplemental Table 2). ADNI had 1205 patients and 6066 visits while NACC had 8121 patients and 35,423 visits. The same preprocessing steps discussed in the previous section were applied to prepare ADNI and NACC datasets.

#### 2.1.3 Dataset notations

After preprocessing, we prepared the datasets as a multivariate longitudinal data. Let *M* denote a dataset with *N* samples (patients), *M* = (*X*_1_, …, *X*_*N*_) where each sample *X* represents measurements of F features collected over *T* time points (visits): *X*= {*x*_1_, *x*_2_, . . ., *x*_*T*_} ∈ ℝ^*T*×*F*^. For each visit *t* = 1,2, . . ., *T, x*_*t*_ = 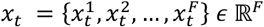 represents a vector of features of sample X at visit *t*. For each feature *f* = 1,2, . . ., *F*, 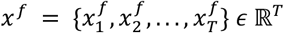 represents the *f*^th^ feature value of sample *X*over *T* visits, and 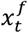represents the *f*^th^ feature value of sample *X*at visit *t*. In *M*, each sample has a corresponding diagnosis (*Y*_1_, …, *Y*_*N*_) for each time point: *Y* = {*y*_1_, *y*_2_, . . ., *y*_*T*_ } ∈ ℝ^*T*×1^. For each visit *t* ∈ {1,2, . . ., *T*}, *y*_*t*_ ∈ {0, 1} where 0 denotes MCI and 1 denotes AD.

## 2.2 Method

In this study, we built two RNN-based deep learning models to predict conversion from MCI to AD at the next visit and the multiple visits ahead. In both models, we utilized the age feature to alleviate the limitation of RNN models with irregular time intervals between consecutive inputs in the sequence. Our proposed models also utilized a customized binary cross entropy loss function to give a higher weight to its sensitivity since early prediction of conversion cases correctly is more important than making a false positive prediction. In the proposed models, the type RNN cell was a hyperparameter with possible choices of LSTM, GRU, Bidirectional LSTM (Bi-LSTM), and Bidirectional GRU (Bi-GRU).

### 2.2.1 A Primer on GRU and Bi-GRU

In GRU, each cell consists of reset (Eq 1) and update gate (Eq 2) though which the cell determines what portion of the previous cell state and the current input will be utilized. To compute the hidden state at time point *t, h*_*t*_, first a candidate hidden state, 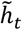 is computed (Eq 3) by utilizing the current input vector *x*_*t*_, the reset gate *r*_*t*_, and the hidden state at the previous time point, *h*_*t*−1_. Then, utilizing the update gate *z*_*t*_, the current hidden state is computed as a weighted average of *h*_*t*−1_ and 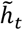 (Eq 4). In Eq 1-4, *W*_*r*_, *U*_*r*_, *W*_*z*_, *U*_*z*_, *W*_*h*_, and *U*_*h*_ are the trainable linear transformation matrices; b_*r*_, b_*z*_, and b_*h*_ are the bias vectors; σ is the sigmoid function and ⊙ is the Hadamard product.

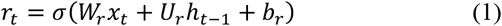

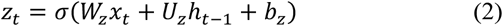

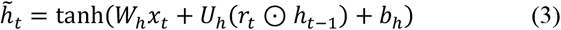

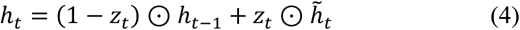

In Bi-GRU, two unidirectional GRUs are used to learn information from previous and later inputs in the sequence while processing the current input (Liu et al., 2021). The first GRU is a forward GRU (*GRU*_*f*_), which is explained in the previous paragraph. The second GRU is a backward GRU (*GRU*_b_) which is exactly same to (*GRU*_*f*_) except that the hidden state of the cell is computed based on the current and later inputs. In other words, the hidden state of a backward GRU cell is calculated based on Eq 1-4 except that all *h*_*t*−1_ terms are replaced with *h*_*t*+1_. To compute the hidden state of Bi-GRU at time point *t*, the hidden states of *GRU*_*f*_ and *GRU*_b_ are computed and concatenated (Eq 5). In Eq 5, ⊕ denotes the concatenation operation.

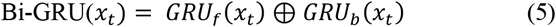

### 2.2.2 Prediction model for conversion to AD at the next visit

To predict AD conversion at the next visit, we developed a framework named Predicting Progression of Alzheimer’s Disease (PPAD) that consists of a RNN and multi-layer perceptron (MLP) (Figure 1). In this architecture, the RNN component learns 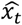, a latent representation of the longitudinal clinical data up to *t* visits (Eq 6). Then, the MLP model is trained with concatenation of the cross-sectional demographic data (*D*) and 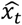 to predict conversion to AD at next visit (Eq 7).

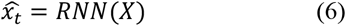

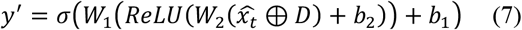

**Fig. 1.**
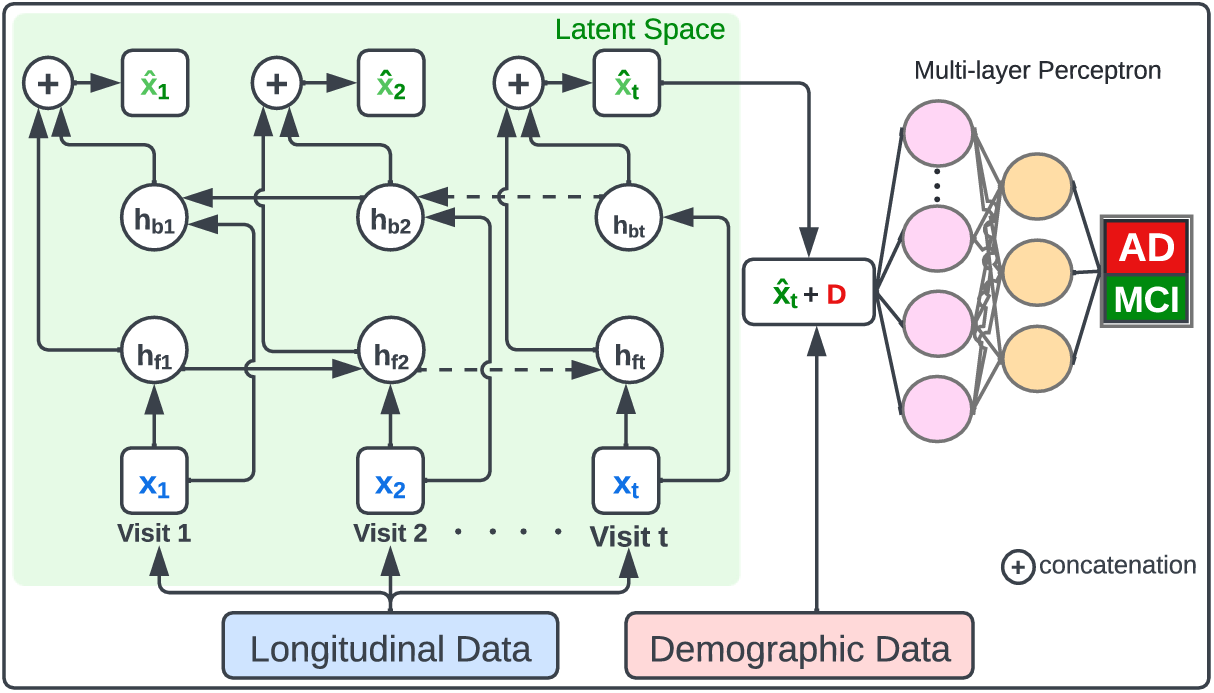
The architecture of PPAD to predict the conversion to AD at the next visit

In Eq 7, *y*′ represents the predicted diagnosis, *W*_1_ and *W*_2_ are the trainable linear transformation matrices, and b_1_ and b_2_ are the bias vectors.

### 2.2.3 Prediction model for conversion to AD at multiple visits ahead

For early predicting conversion of AD at multiple visits ahead, we propose another architecture PPAD-AE that composes of a RNN autoencoder and an MLP (Figure 2). In this architecture, the RNN component learns a latent representation 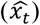 of the longitudinal clinical data up to *t* visits (Eq 6). Then, the latent representation is used by the decoder component to generate representations of multiple visits ahead up to *n* visits. Finally, the MLP model is trained with concatenation of the cross-sectional demographic data (*D*) and the representation of the last generated visit by the decoder to predict conversion to AD at the (*t* + *n*)^*th*^ visit (Eq 8).

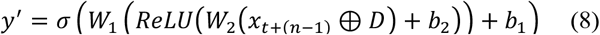

**Fig. 2.**
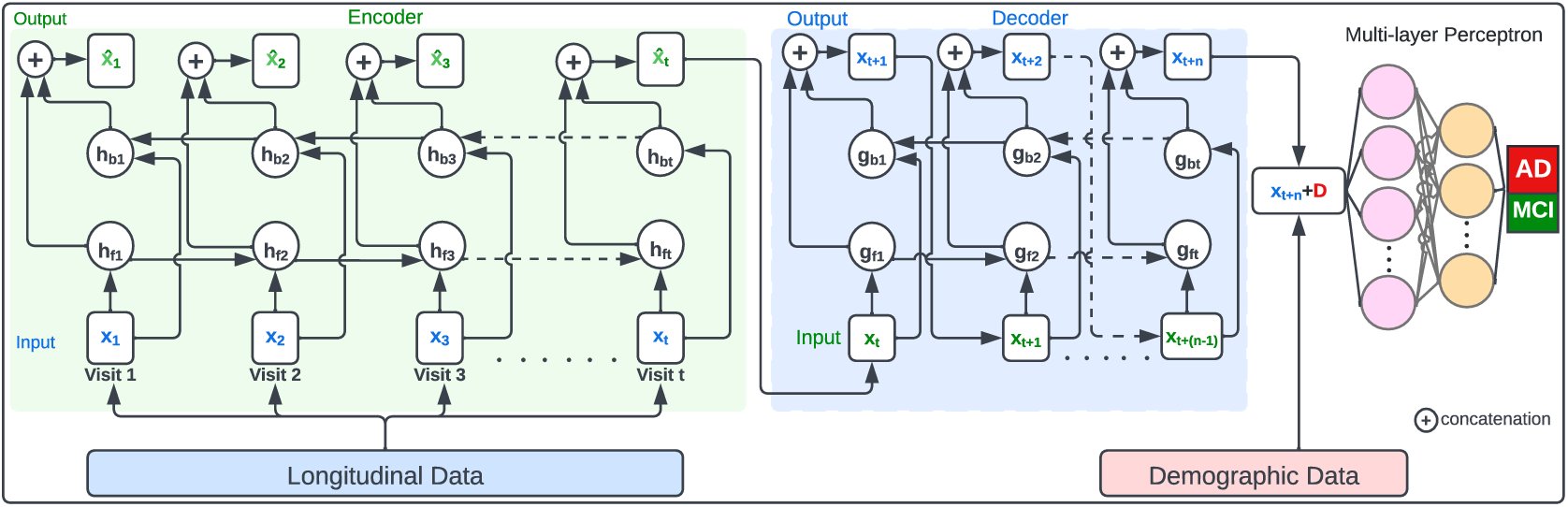
The architecture of PPAD-AE to predict the conversion to AD at the future visits.

### 2.2.4 Parameter learning and evaluation metrics

To increase the prediction’s sensitivity for both architectures, all trainable parameters for the RNN, RNN autoencoder, and MLP were learned in an integral way using a customized binary cross-entropy loss function (Eq 9) to give more weight on predicting conversion to AD cases. We seek by using this customized loss function to minimize the false negative cases while predicting diagnosis of the future visit which leads to increased sensitivity of the predictive model.

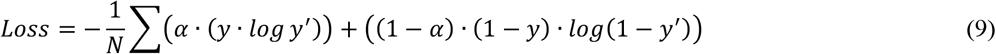

In Eq 9, α is a real number between 0 and 1 to define the relative weight of positive prediction, *y* is the true diagnosis, and *y*′ is the predicted diagnosis. In this study, we set α to 0.7. Based on the proposed customized loss function, all trainable parameters for the RNN, RNN autoencoder, and MLP (Eq 7 and 8) were updated while training models in the backpropagation. For optimization, all models were trained using Adaptive Moment Estimation (Adam) optimizer and the learning rate was set to 0.001. RNN cell, number of epochs, batch size, dropout rate, and L2 regularization are the hyperparameters that have been tuned. For model evaluation, F2 score (Eq 10) and sensitivity were used.

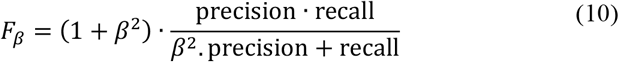

In Eq 10, recall is considered β times more important than precision. In this study, β was set to 2.

## 3 Results and Discussion

In this study, we propose two RNN-based architectures, namely PPAD and PPAD-AE for the prediction of conversion to AD at the next visit and the multiple visits ahead, respectively. We evaluated the proposed architectures in two experimental setups. In the first setup, ADNI dataset was utilized to train and test the proposed architectures using the longitudinal multi-modal and the cross-sectional demographic data. The longitudinal data consisted of 20 features from cognitive performance and neuroimaging biomarker data modalities (Supplemental Table 1). The cross-sectional demographic data consisted of gender, race, ethnicity, education, and APOEɛ4. We split data to 70% training and 30% test three times and reported the average performance across these splits. In the second setup, the models were trained on the entire ADNI longitudinal and cross-sectional data and tested on NACC data. The longitudinal data consisted of five features and the cross-sectional demographic data consisted of gender, race, ethnicity, education, and APOEɛ4 (Supplemental Table 2). For both experimental setups, we used age as a longitudinal feature to represent the time difference between consecutive visits. To select the optimal values for the hyperparameters (i.e., RNN cell, number of epochs, batch size, dropout rate, and L2 regularization), we performed grid search with 5-fold cross validation for each investigated scenario in both experimental setups. Supplemental Table 3 and 4 show the best hyperparameter values for PPAD and PPAD-AE for all splits in the first experimental setup, respectively. Supplemental Table 5 and 6 show the best hyperparameter values for PPAD and PPAD-AE in the second experimental setup, respectively. Supplemental Table 7 and 8 show the number of converters and non-converters in each scenario in the first and second experimental setup, respectively.

### 3.1 Predicting the conversion to AD at the next visit

To evaluate PPAD, which predicts conversion to AD at the next visit, we trained it using different scenarios for both experimental setups. Specifically, we trained four models using two, three, five, and six visits to predict the conversion to AD at the next visit (i.e., at the third, fourth, sixth and seventh visit, respectively). For comparison, we trained a TLSTM-based architecture using the same training data. Since T-LSTM can handle irregular intervals between visits internally, to check the effectiveness of utilizing the age feature, we did not use the age feature for T-LSTM. In addition, Random Forest- (RF) and SVM-based models were trained as baseline. Since RF and SVM cannot handle longitudinal data, they were trained using the same training data used for RNN models after aggregating each feature value by computing its mean. For generalization, the whole procedure was repeated 15 times. The results on both experimental setups showed that PPAD outperformed all baseline models in terms of F2 (Figure 3A and 3B) and sensitivity (Supplemental Figure 1A and 1B). The results also showed that, as expected, training using more visits improved the performance of next visit diagnosis prediction for all RNN models. In addition, PPAD outperformed T-LSTM in seven out of eight cases, which shows the ability of our model to address the irregular time intervals problem better than T-LSTM.

**Fig. 3.**
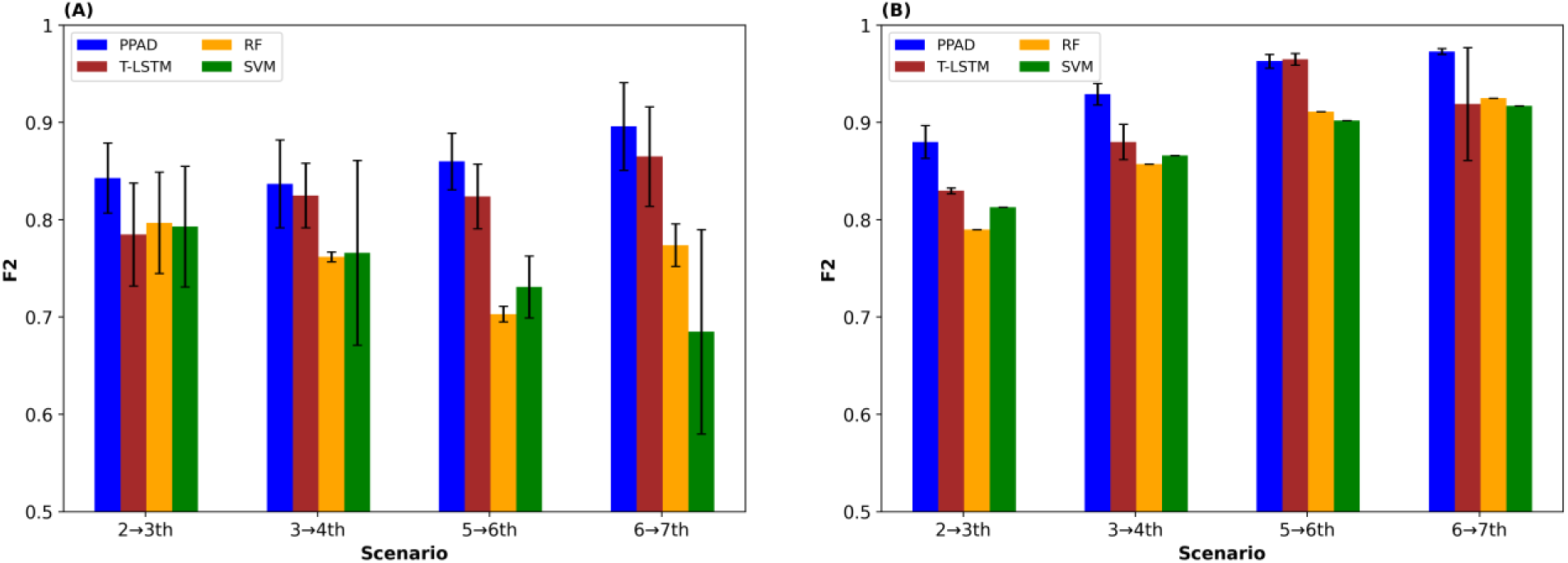
F2 scores for PPAD models to predict conversion to AD at the next visit. (A) Models tested on held-out samples in ADNI after training using 2, 3, 5, and 6 visits in ADNI, respectively. (B) Models tested on NACC after training using 2, 3, 5, and 6 visits in ADNI, respectively.

### 3.2 Predicting the conversion to AD at multiple visits ahead

To evaluate PPAD-AE, which predicts the conversion to AD at multiple visits ahead, we trained it using different scenarios. Specifically, we trained four models using two, three, five and six visits to predict the diagnosis at the following second, third, and fourth visits ahead. For example, the model that was trained using two visits was evaluated on predicting the diagnosis at the fourth, fifth and sixth visits. For generalization, the whole procedure was repeated 15 times. We compared PPAD-AE to RF and SVM. T-LSMT was unable to predict multiple visits ahead, thus was not used in this evaluation. We observed that PPAD-AE outperformed all baseline models in terms of F2 (Figure 4) and sensitivity (Supplemental Figure 2) in both experimental setups except for one scenario (Figure 4F and Supplemental Figure 2F). As observed in the PPAD results (Figure 3), training the model with more visits improved the prediction performance in general, whereas the performance of most models dropped when predicting diagnosis at the farther time points. We also observed that the performance of PPAD-AE on NACC dataset was higher than held-out dataset of ADNI. This could be due to using a larger training set (i.e., the entire ADNI data) when testing the model on NACC cohort.

**Fig. 4.**
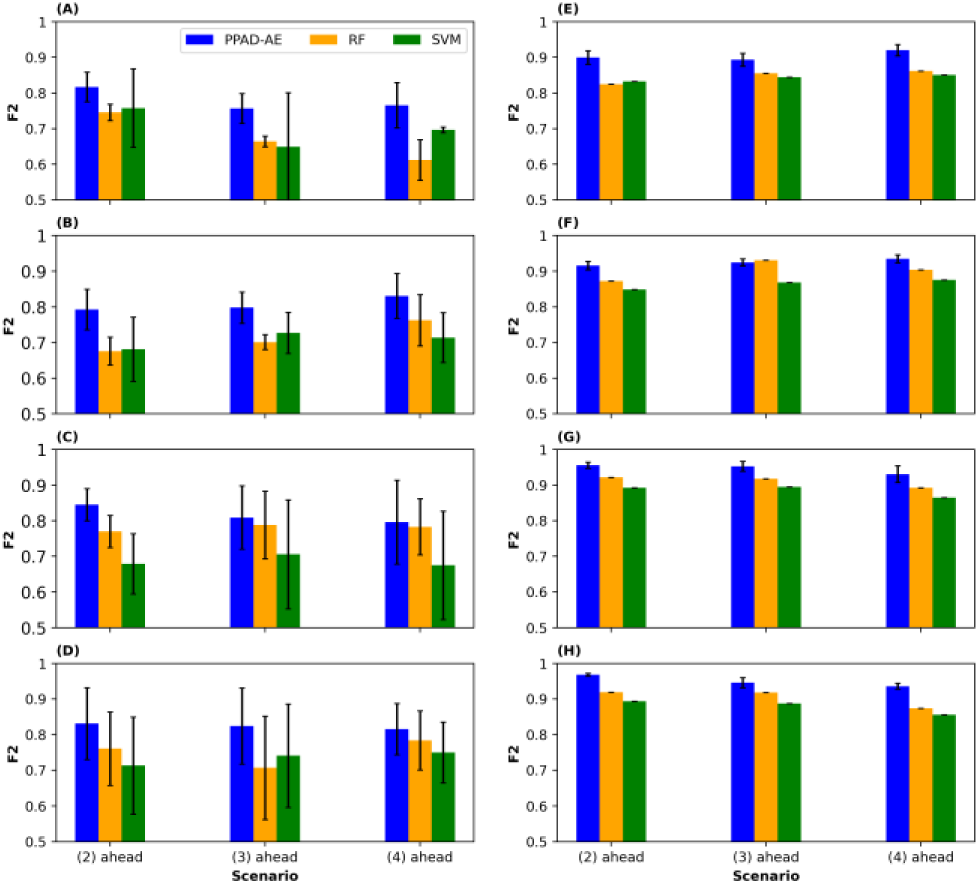
F2 scores for PPAD-AE models to predict conversion to AD at the second, third, and fourth visits ahead. (A, B, C, and D) Models tested on held-out samples in ADNI after training using 2, 3, 5, and 6 visits in ADNI, respectively. (E, F, G, and H) Models tested on NACC after training using 2, 3, 5, and 6 visits in ADNI, respectively.

### 3.3 Feature importance Analysis

We investigated the performance of the proposed models (Figure 1 and 2) to determine feature importance through SHapley Additive exPlanations (SHAP). Figure 5 and Supplemental Figure 3 show the mean absolute SHAP value for the longitudinal features for the proposed models with first and second experimental setups, respectively. The results indicate that Functional Activities Questionnaire (FAQ) and the logical memory delayed recall total (LDELTOTAL) are the most important features to predict conversion to AD in all scenarios. In addition, age is among important features in predicting conversion to AD. For example, age was the seventh important feature among 20 features (Figure 5A).

**Fig. 5.**
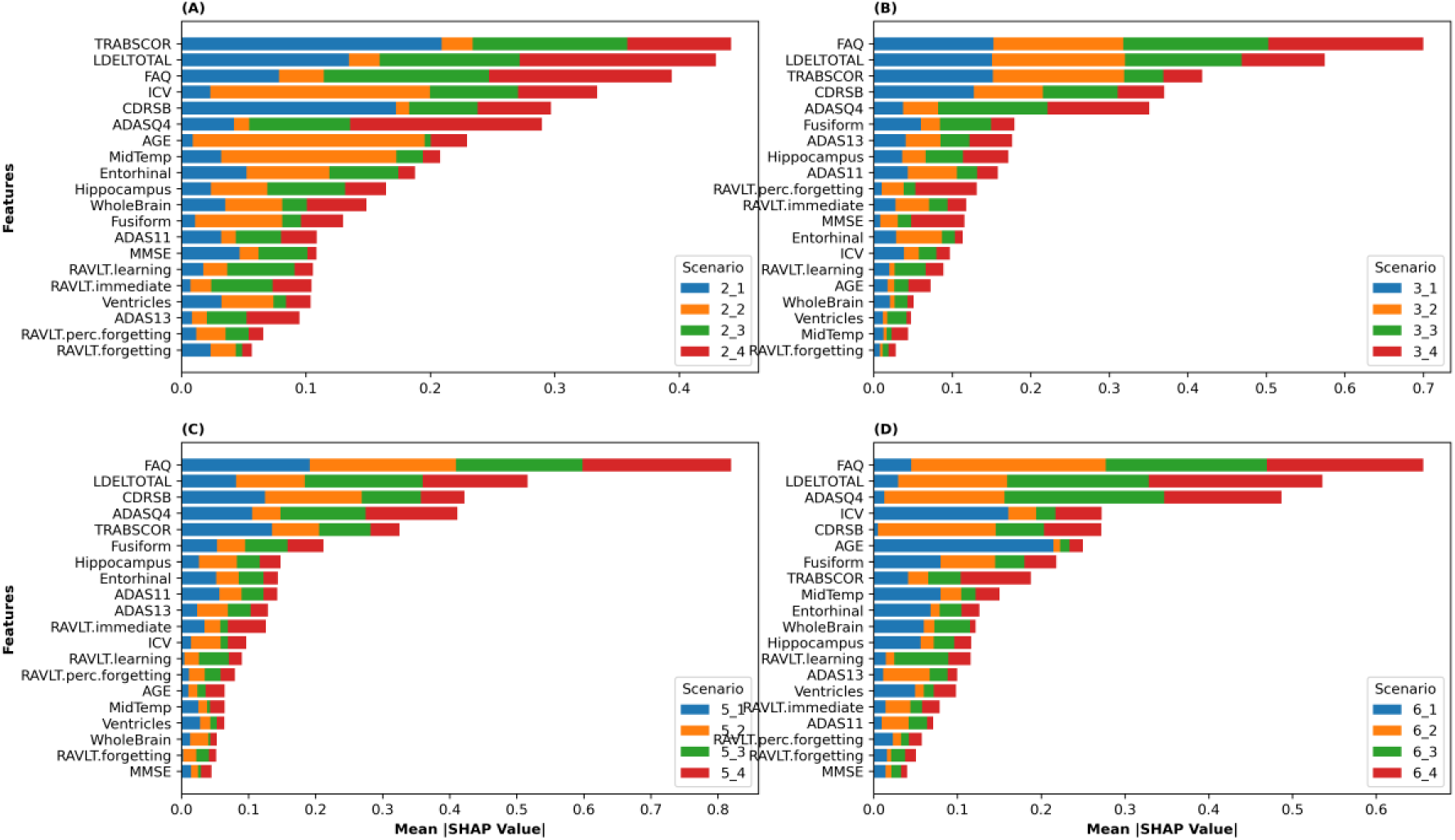
SHAP values for all features used in the first experimental setup using A) 2 B) 3 C) 5 D) 6 visits to train the models. 2_1 means trained using two visits to predict conversion to AD at the next visit, 3_2 means trained using three visits to predict conversion at two visits ahead, and so on.

## 4 Conclusion

In this study, we present two deep learning architectures, PPAD and PPAD-AE to predict the conversion to AD in the future. We utilized longitudinal cognitive and neuroimaging features and cross-sectional demographic data from two large AD databases to evaluate our models. We conducted two experimental setups where ADNI data was used partially or completely to train the model and held-out data and NACC data were used to test the models. In both experimental setups, our tools outperformed other existing tools and baseline models. We also investigated other tools, but could not test them as the code or the tool was not made available.

By utilizing a customized loss function, we gave higher emphasis on the sensitivity of the models. Because for AD prediction, false positive (predicting someone to convert to AD falsely) is less severe than a false negative (not being able to predict a conversion case). PPAD and PPAD-AE are also flexible to incorporate additional features including omics features from gene expression, DNA methylation datasets and blood-based biomarker measurements. To increase its usability, we make PPAD and PPAD-AE publicly available at https://github.com/bozdaglab/PPAD/.

## Acknowledgements

Data collection and sharing for this project was funded by the Alzheimer’s Disease Neuroimaging Initiative (ADNI) (National Institutes of Health Grant U01 AG024904) and DOD ADNI (Department of Defense award number W81XWH-12-2-0012). ADNI is funded by the National Institute on Aging, the National Institute of Biomedical Imaging and Bioengineering, and through generous contributions from the following: AbbVie, Alzheimer’s Association; Alzheimer’s Drug Discovery Foundation; Araclon Biotech; BioClinica, Inc.; Biogen; Bristol-Myers Squibb Company; CereSpir, Inc.; Cogstate; Eisai Inc.; Elan Pharmaceuticals, Inc.; Eli Lilly and Company; EuroImmun; F. Hoffmann-La Roche Ltd and its affiliated company Genentech, Inc.; Fujirebio; GE Healthcare; IXICO Ltd.; Janssen Alzheimer Immunotherapy Research & Development, LLC.; Johnson & Johnson Pharmaceutical Research & Development LLC.; Lumosity; Lundbeck; Merck & Co., Inc.; Meso Scale Diagnostics, LLC.; NeuroRx Research; Neurotrack Technologies; Novartis Pharmaceuticals Corporation; Pfizer Inc.; Piramal Imaging; Servier; Takeda Pharmaceutical Company; and Transition Therapeutics. The Canadian Institutes of Health Research is providing funds to support ADNI clinical sites in Canada. Private sector contributions are facilitated by the Foundation for the National Institutes of Health (www.fnih.org). The grantee organization is the Northern California Institute for Research and Education, and the study is coordinated by the Alzheimer’s Therapeutic Research Institute at the University of Southern California. ADNI data are disseminated by the Laboratory for Neuro Imaging at the University of Southern California.

The NACC database is funded by NIA/NIH Grant U24 AG072122. NACC data are contributed by the NIA-funded ADRCs: P30 AG062429 (PI James Brewer, MD, PhD), P30 AG066468 (PI Oscar Lopez, MD), P30 AG062421 (PI Bradley Hyman, MD, PhD), P30 AG066509 (PI Thomas Grabowski, MD), P30 AG066514 (PI Mary Sano, PhD), P30 AG066530 (PI Helena Chui, MD), P30 AG066507 (PI Marilyn Albert, PhD), P30 AG066444 (PI John Morris, MD), P30 AG066518 (PI Jeffrey Kaye, MD), P30 AG066512 (PI Thomas Wisniewski, MD), P30 AG066462 (PI Scott Small, MD), P30 AG072979 (PI David Wolk, MD), P30 AG072972 (PI Charles DeCarli, MD), P30 AG072976 (PI Andrew Saykin, PsyD), P30 AG072975 (PI David Bennett, MD), P30 AG072978 (PI Neil Kowall, MD), P30 AG072977 (PI Robert Vassar, PhD), P30 AG066519 (PI Frank LaFerla, PhD), P30 AG062677 (PI Ronald Petersen, MD, PhD), P30 AG079280 (PI Eric Reiman, MD), P30 AG062422 (PI Gil Rabinovici, MD), P30 AG066511 (PI Allan Levey, MD, PhD), P30 AG072946 (PI Linda Van Eldik, PhD), P30 AG062715 (PI Sanjay Asthana, MD, FRCP), P30 AG072973 (PI Russell Swerdlow, MD), P30 AG066506 (PI Todd Golde, MD, PhD), P30 AG066508 (PI Stephen Strittmatter, MD, PhD), P30 AG066515 (PI Victor Henderson, MD, MS), P30 AG072947 (PI Suzanne Craft, PhD), P30 AG072931 (PI Henry Paulson, MD, PhD), P30 AG066546 (PI Sudha Seshadri, MD), P20 AG068024 (PI Erik Roberson, MD, PhD), P20 AG068053 (PI Justin Miller, PhD), P20 AG068077 (PI Gary Rosenberg, MD), P20 AG068082 (PI Angela Jefferson, PhD), P30 AG072958 (PI Heather Whitson, MD), P30 AG072959 (PI James Leverenz, MD).

## Funding

This work was supported by the National Institute of General Medical Sciences of the National Institutes of Health under Award Number R35GM133657 and the startup funds from the University of North Texas.

## Notes

### Competing Interest Statement

The authors have declared no competing interest.

### Summary of Updates

Figure 5 was updated

