## Supplemental Table for "PPAD: A deep learning architecture to predict progression of Alzheimer’s disease" (from another version of this preprint: version 1, 2023-01-30)

Supplemental Table 1 List of features in ADNI data used to train the models in the first experimental setup where models were trained on 70% of ADNI data and tested on the 30% held-out ADNI data. MRI: Magnetic Resonance Imaging.

| Feature name | Description | Category | Type |
|---|---|---|---|
| **CDRSB** | Clinical Dementia Rating Scale–Sum of Boxes | Cognitive performance | Longitudinal |
| **ADAS11** | Alzheimer's Disease Assessment Scale 11 | Cognitive performance | Longitudinal |
| **ADAS13** | Alzheimer's Disease Assessment Scale 13 | Cognitive performance | Longitudinal |
| **ADASQ4** | Alzheimer's Disease Assessment Scale Q4 | Cognitive performance | Longitudinal |
| **MMSE** | Mini-Mental State Examination | Cognitive performance | Longitudinal |
| **RAVLT.immediate** | Rey Auditory Verbal Learning Test (Immediate recall) | Cognitive performance | Longitudinal |
| **RAVLT.learning** | Rey Auditory Verbal Learning Test (Learning) | Cognitive performance | Longitudinal |
| **RAVLT.forgetting** | Rey Auditory Verbal Learning Test (Forgetting) | Cognitive performance | Longitudinal |
| **RAVLT.perc.forgetting** | Rey Auditory Verbal Learning Test (Percent Forgetting) | Cognitive performance | Longitudinal |
| **LDELTOTAL** | The logical memory delayed recall total | Cognitive performance | Longitudinal |
| **TRABSCOR** | Trail Making Test-B | Cognitive performance | Longitudinal |
| **FAQ** | Functional Activities Questionnaire | Cognitive performance | Longitudinal |
| **Ventricles** | Ventricles measurement | MRI | Longitudinal |
| **Hippocampus** | Hippocampus measurement | MRI | Longitudinal |
| **WholeBrain** | Whole Brain measurement | MRI | Longitudinal |
| **Entorhinal** | Entorhinal cortex measurement | MRI | Longitudinal |
| **Fusiform** | Fusiform Gyrus | MRI | Longitudinal |
| **MidTemp** | Scan of the middle temporal artery | MRI | Longitudinal |
| **ICV** | Intracranial volume | MRI | Longitudinal |
| **AGE** | Age of patient | Demographic | Longitudinal |
| **PTGENDER** | Gender of patient | Demographic | Cross-sectional |
| **PTEDUCAT** | Education years of patient | Demographic | Cross-sectional |
| **PTETHCAT** | Ethnicity of patient | Demographic | Cross-sectional |
| **PTRACCAT** | Race of patient | Demographic | Cross-sectional |
| **APOE4** | Number of ε4 allele | Demographic | Cross-sectional |

Supplemental Table 2 List of features in NACC data for the second setup, where models were trained on ADNI data and tested on NACC data.

| Feature name | Corresponding feature in ADNI |
|---|---|
| **CDRSUM** | CDRSB |
| **NACCMMSE** | MMSE |
| **MEMUNITS** | LDELTOTAL |
| **FAQ*** | FAQ |
| **NACCAGE** | AGE |
| **SEX** | PTGENDER |
| **EDUC** | PTEDUCAT |
| **HISPANIC** | PTETHCAT |
| **RACE** | PTRACCAT |
| **NACCNE4S** | APOE4 |

* NACC dataset does not have FAQ feature, but it has the raw data that can be used to get it based on the following equation:

FAQ = BILLS+ TAXES+ SHOPPING+ GAMES+STOVE+MEALPREP+EVENTS+PAYATTN+ REMDATES+TRAVEL

The value of each feature that contributes FAQ value has the following range (0 = Normal, 1 = Has difficulty but does by self, 2 = Requires assistance, 3 = Dependent).

Supplemental Table 3 Optimal hyperparameter values of PPAD for each scenario in all splits in the first experimental setup. For each scenario, $x \rightarrow 1$ means that the model was trained using $x$ visits and tested the diagnosis at the next visit.

| Data split | Scenario | RNN cell | Batch size | Epochs | Dropout rate | L2 |
|---|---|---|---|---|---|---|
| **1** | 2→1 | Bi-GRU | 4 | 50 | 0.4 | $1e^{-5}$ |
| | 3→1 | Bi-GRU | 16 | 20 | 0.2 | $1e^{-3}$ |
| | 5→1 | Bi-LSTM | 2 | 50 | 0.1 | $1e^{-5}$ |
| | 6→1 | Bi-GRU | 8 | 40 | 0.2 | $1e^{-5}$ |
| **2** | 2→1 | GRU | 16 | 100 | 0.4 | $1e^{-3}$ |
| | 3→1 | GRU | 2 | 40 | 0.0 | $1e^{-3}$ |
| | 5→1 | Bi-GRU | 4 | 20 | 0.0 | $1e^{-3}$ |
| | 6→1 | Bi-GRU | 16 | 30 | 0.3 | $1e^{-7}$ |

| | | | | | | |
|---|---|---|---|---|---|---|
| **3** | 2→1 | Bi-LSTM | 32 | 30 | 0.0 | $1e^{-5}$ |
| | 3→1 | Bi-GRU | 4 | 50 | 0.2 | $1e^{-3}$ |
| | 5→1 | Bi-LSTM | 8 | 50 | 0.5 | $1e^{-7}$ |
| | 6→1 | LSTM | 2 | 40 | 0.4 | $1e^{-3}$ |

Supplemental Table 4 Optimal hyperparameter values of PPAD-AE for each scenario in all splits in the first experimental setup. For each scenario, $x \rightarrow y$ means that the model was trained using $x$ visits and tested the diagnosis at the next $y^{th}$ visit ahead.

| Data split | Scenario | RNN cell | Batch size | Epochs | Dropout rate | L2 |
|---|---|---|---|---|---|---|
| **1** | 2→1 | Bi-LSTM | 8 | 40 | 0.1 | 1e-3 |
| | 2→2 | GRU | 4 | 10 | 0.1 | 1e-3 |
| | 2→3 | Bi-GRU | 16 | 30 | 0.1 | 1e-3 |
| | 2→4 | Bi-GRU | 4 | 20 | 0.0 | 1e-3 |
| | 3→1 | Bi-GRU | 4 | 10 | 0.1 | 1e-3 |
| | 3→2 | Bi-GRU | 2 | 40 | 0.5 | 1e-5 |
| | 3→3 | LSTM | 4 | 40 | 0.4 | 1e-5 |
| | 3→4 | GRU | 8 | 30 | 0.0 | 1e-3 |
| | 5→1 | GRU | 4 | 50 | 0.1 | 1e-7 |
| | 5→2 | Bi-LSTM | 2 | 20 | 0.0 | 1e-3 |
| | 5→3 | GRU | 2 | 40 | 0.3 | 1e-7 |
| | 5→4 | Bi-GRU | 2 | 30 | 0.0 | 1e-5 |
| | 6→1 | GRU | 4 | 30 | 0.0 | 1e-5 |
| | 6→2 | Bi-GRU | 8 | 30 | 0.1 | 1e-5 |
| | 6→3 | LSTM | 2 | 10 | 0.1 | 1e-3 |
| | 6→4 | Bi-GRU | 4 | 40 | 0.1 | 1e-3 |
| **2** | 2→1 | Bi-LSTM | 4 | 10 | 0.4 | 1e-7 |
| | 2→2 | Bi-GRU | 8 | 20 | 0.1 | 1e-3 |
| | 2→3 | Bi-LSTM | 2 | 30 | 0.1 | 1e-7 |
| | 2→4 | Bi-GRU | 32 | 20 | 0.0 | 1e-3 |
| | 3→1 | GRU | 32 | 100 | 0.3 | 1e-7 |
| | 3→2 | Bi-GRU | 32 | 20 | 0.0 | 1e-5 |

| | | | | | | |
|---|---|---|---|---|---|---|
| | 3→3 | Bi-GRU | 8 | 40 | 0.2 | 1e-7 |
| | 3→4 | GRU | 2 | 30 | 0.3 | 1e-5 |
| | 5→1 | Bi-LSTM | 8 | 100 | 0.4 | 1e-5 |
| | 5→2 | Bi-LSTM | 32 | 100 | 0.2 | 1e-7 |
| | 5→3 | Bi-GRU | 16 | 20 | 0.0 | 1e-5 |
| | 5→4 | GRU | 2 | 50 | 0.2 | 1e-3 |
| | 6→1 | Bi-GRU | 2 | 20 | 0.0 | 1e-7 |
| | 6→2 | Bi-GRU | 2 | 100 | 0.4 | 1e-5 |
| | 6→3 | LSTM | 2 | 30 | 0.2 | 1e-7 |
| | 6→4 | Bi-GRU | 2 | 40 | 0.1 | 1e-5 |
| **3** | 2→1 | Bi-LSTM | 4 | 10 | 0.3 | 1e-5 |
| | 2→2 | GRU | 32 | 50 | 0.1 | 1e-5 |
| | 2→3 | GRU | 4 | 40 | 0.2 | 1e-5 |
| | 2→4 | Bi-GRU | 16 | 10 | 0.1 | 1e-3 |
| | 3→1 | Bi-GRU | 4 | 100 | 0.4 | 1e-5 |
| | 3→2 | GRU | 8 | 30 | 0.1 | 1e-7 |
| | 3→3 | Bi-GRU | 16 | 30 | 0.1 | 1e-7 |
| | 3→4 | Bi-GRU | 8 | 20 | 0.4 | 1e-3 |
| | 5→1 | Bi-GRU | 4 | 50 | 0.4 | 1e-7 |
| | 5→2 | Bi-GRU | 8 | 50 | 0.4 | 1e-7 |
| | 5→3 | GRU | 4 | 10 | 0.1 | 1e-7 |
| | 5→4 | Bi-GRU | 2 | 40 | 0.5 | 1e-5 |
| | 6→1 | Bi-GRU | 2 | 100 | 0.4 | 1e-5 |
| | 6→2 | Bi-LSTM | 4 | 20 | 0.2 | 1e-5 |
| | 6→3 | Bi-LSTM | 2 | 20 | 0.4 | 1e-7 |
| | 6→4 | Bi-GRU | 16 | 30 | 0.2 | 1e-5 |

Supplemental Table 5 Optimal hyperparameter values of PPAD for each scenario in all splits in the second experimental setup. For each scenario, $x \rightarrow 1$ means that the model was trained using $x$ visits and tested the diagnosis at the next visit.

| Scenario | RNN cell | Batch size | Epochs | Dropout rate | L2 |
|---|---|---|---|---|---|
| **2→1** | GRU | 2 | 40 | 0.1 | $1e^{-7}$ |
| **3→1** | Bi-GRU | 32 | 20 | 0.2 | $1e^{-3}$ |
| **5→1** | Bi-LSTM | 4 | 10 | 0.0 | $1e^{-7}$ |
| **6→1** | Bi-GRU | 4 | 30 | 0.2 | $1e^{-5}$ |

Supplemental Table 6 Optimal hyperparameter values of PPAD-AE for each scenario in all splits in the second experimental setup. For each scenario, $x \rightarrow y$ means that the model was trained using $x$ visits and tested the diagnosis at the next $y^{th}$ visit ahead.

| Scenario | RNN cell | Batch size | Epochs | Dropout rate | L2 |
|---|---|---|---|---|---|
| **2→1** | GRU | 2 | 40 | 0.2 | 1e-7 |
| **2→2** | Bi-GRU | 4 | 20 | 0.2 | 1e-7 |
| **2→3** | GRU | 16 | 10 | 0.0 | 1e-7 |
| **2→4** | Bi-GRU | 8 | 10 | 0.0 | 1e-7 |
| **3→1** | Bi-GRU | 2 | 10 | 0.4 | 1e-3 |
| **3→2** | Bi-GRU | 16 | 20 | 0.1 | 1e-5 |
| **3→3** | GRU | 8 | 30 | 0.1 | 1e-7 |
| **3→4** | GRU | 16 | 40 | 0.0 | 1e-3 |
| **5→1** | Bi-LSTM | 2 | 30 | 0.3 | 1e-5 |
| **5→2** | Bi-GRU | 2 | 10 | 0.2 | 1e-3 |
| **5→3** | Bi-LSTM | 4 | 30 | 0.3 | 1e-5 |
| **5→4** | Bi-GRU | 16 | 10 | 0.0 | 1e-5 |
| **6→1** | Bi-LSTM | 4 | 30 | 0.1 | 1e-7 |
| **6→2** | GRU | 64 | 100 | 0.1 | 1e-7 |
| **6→3** | GRU | 8 | 10 | 0.1 | 1e-5 |
| **6→4** | GRU | 8 | 50 | 0.4 | 1e-3 |

Supplemental Table 7 Summary of number of Non-converter and converter cases in each scenario in the first experimental setup

| | **Train** | | **Test** | |
|---|---|---|---|---|
| **Scenario** | **Non-Converter** | **Converter** | **Non-Converter** | **Converter** |
| **2_1** | 499 | 275 | 211 | 132 |
| **2_2** | 378 | 294 | 159 | 108 |
| **2_3** | 291 | 231 | 124 | 61 |
| **2_4** | 219 | 140 | 84 | 53 |
| **3_1** | 438 | 336 | 159 | 108 |
| **3_2** | 349 | 323 | 124 | 61 |
| **3_3** | 261 | 261 | 84 | 53 |
| **3_4** | 203 | 156 | 47 | 40 |
| **5_1** | 379 | 395 | 84 | 53 |
| **5_2** | 303 | 369 | 47 | 40 |
| **5_3** | 239 | 283 | 32 | 23 |
| **5_4** | 195 | 164 | 23 | 18 |
| **6_1** | 363 | 411 | 47 | 40 |
| **6_2** | 297 | 375 | 32 | 23 |
| **6_3** | 237 | 285 | 23 | 18 |
| **6_4** | 193 | 166 | 17 | 9 |

Supplemental Table 8 Summary of number of Non-converter and converter cases in each scenario in the second experimental setup

| | Train | | Test | |
|---|---|---|---|---|
| **Scenario** | **Non-Converter** | **Converter** | **Non-Converter** | **Converter** |
| **2_1** | 745 | 460 | 1447 | 4671 |
| **2_2** | 569 | 471 | 869 | 3530 |
| **2_3** | 439 | 382 | 522 | 2527 |
| **2_4** | 335 | 233 | 317 | 1800 |
| **3_1** | 666 | 539 | 869 | 3530 |
| **3_2** | 516 | 524 | 522 | 2527 |
| **3_3** | 399 | 422 | 317 | 1800 |
| **3_4** | 307 | 261 | 198 | 1216 |
| **5_1** | 573 | 632 | 317 | 1800 |
| **5_2** | 448 | 592 | 198 | 1216 |
| **5_3** | 361 | 460 | 121 | 768 |
| **5_4** | 292 | 276 | 71 | 480 |
| **6_1** | 545 | 660 | 198 | 1216 |
| **6_2** | 438 | 602 | 121 | 768 |
| **6_3** | 356 | 465 | 71 | 480 |
| **6_4** | 288 | 280 | 44 | 262 |

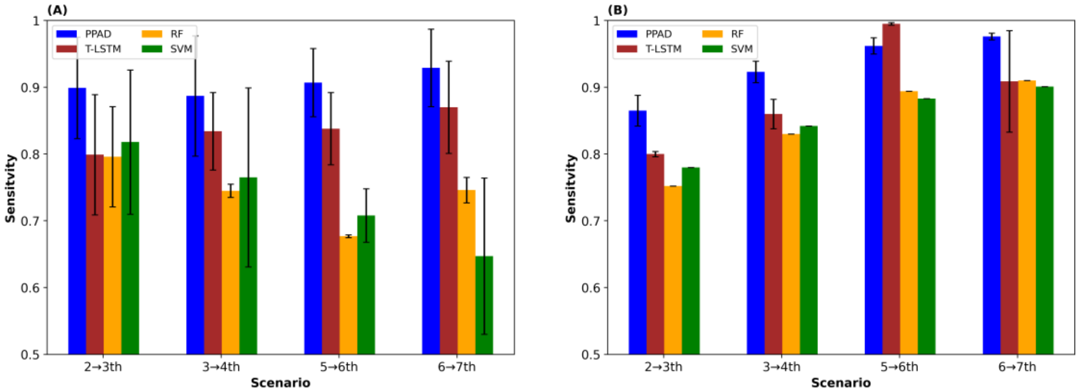


**Supplemental Fig. 1** Sensitivity scores for PPAD models to predict conversion to AD at the next visit. (A) Models tested on held-out samples in ADNI after training using 2, 3, 5, and 6 visits in ADNI, respectively. (B) Models tested on NACC after training using 2, 3, 5, and 6 visits in ADNI, respectively.

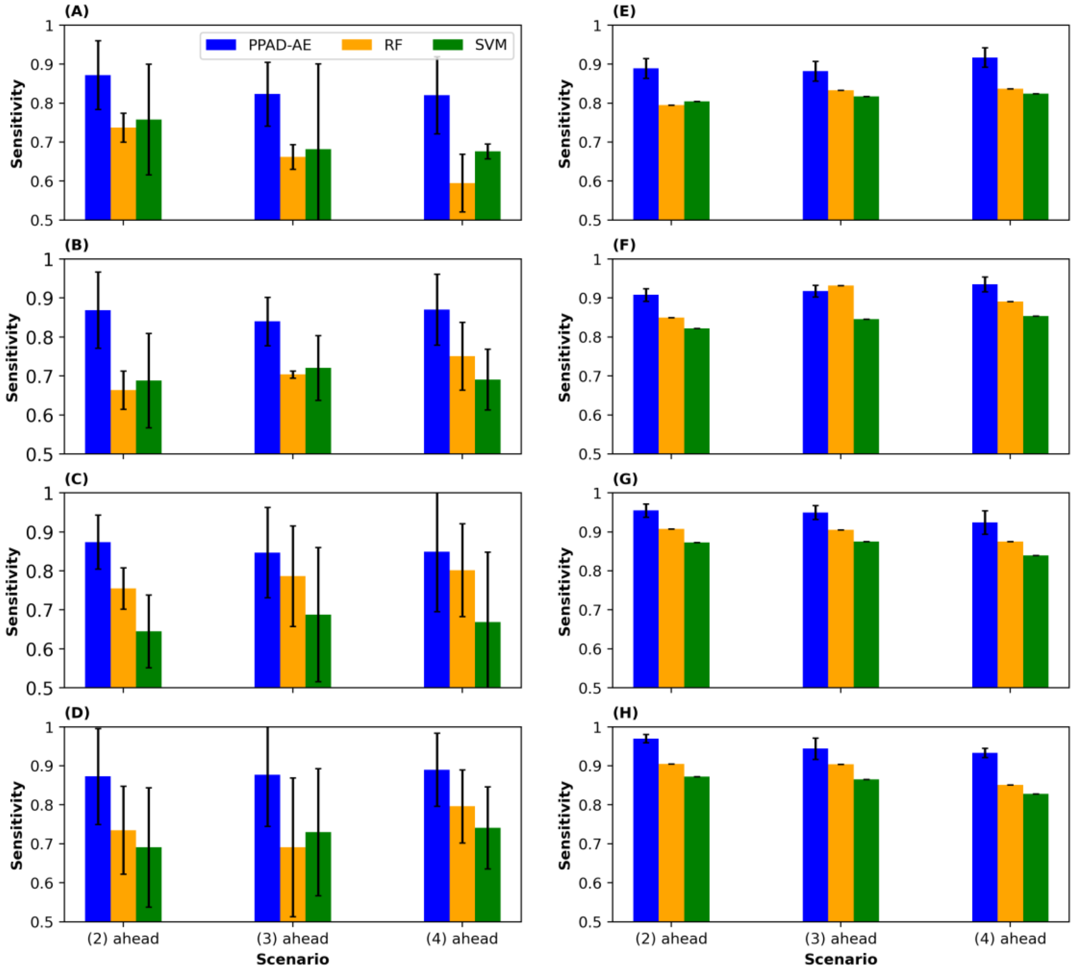


**Supplemental Fig. 2** Sensitivity scores for PPAD-AE models to predict progression to AD at the next second, third, and fourth visits ahead. (A, B, C, and D) Models tested on held-out samples in ADNI after training using 2, 3, 5, and 6 visits in ADNI, respectively. (E, F, G, and H) Models tested on NACC after training using 2, 3, 5, and 6 visits in ADNI, respectively.

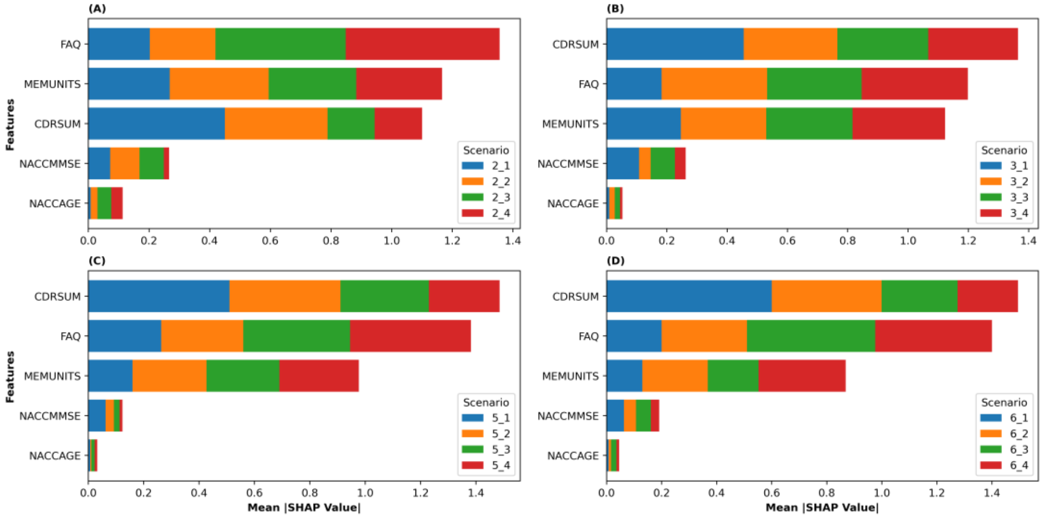


**Supplemental Fig. 3** SHAP values for all features used in the second experimental setup using A) 2 B) 3 C) 5 D) 6 visits to train the models. 2_1 means trained using two visits to predict conversion to AD at the next visit, 3_2 means trained using three visits to predict conversion at two visits ahead, and so on.
